# Does rainforest biodiversity stand on the shoulders of giants? Effect of disturbances by forest elephants on trees and insects on Mount Cameroon

**DOI:** 10.1101/2020.06.10.144279

**Authors:** Vincent Maicher, Sylvain Delabye, Mercy Murkwe, Jiří Doležal, Jan Altman, Ishmeal N. Kobe, Julie Desmist, Eric B. Fokam, Tomasz Pyrcz, Robert Tropek

**Affiliations:** Institute of Entomology, Biology Centre, Czech Academy of Sciences, Branisovska 31, CZ-37005 Ceske Budejovice, Czechia; Faculty of Science, University of South Bohemia, Branisovska 1760, CZ-37005 Ceske Budejovice, Czechia; Nicholas School of the Environment, Duke University, 9 Circuit Dr., Durham, NC 27710, United States of America; Department of Zoology and Animal Physiology, Faculty of Science, University of Buea, P.O. Box 63 Buea, Cameroon; Department of Ecology, Faculty of Science, Charles University, Vinicna 7, CZ-12844 Prague, Czechia; Institute of Botany, Czech Academy of Sciences, Dukelska 135, CZ-37982 Trebon, Czechia; University Paris-Saclay, 15 rue Georges Clemenceau 91400 Orsay, France; Institute of Zoology and Biomedical Research, Jagiellonian University, Gronostajowa 9, PL-30-387 Krakow, Poland; Nature Education Centre of the Jagiellonian University, Gronostajowa 5, PL-30-387 Krakow, Poland

**Keywords:** Afrotropics, Lepidoptera, Megafauna, Megaherbivores, Natural disturbances, Trees

## Abstract

Natural disturbances are essential for dynamics of tropical rainforests, contributing to their tremendous biodiversity. In the Afrotropical rainforests, megaherbivores have played a key role before their recent decline. Although the influence of savanna elephants on ecosystems has been documented, their close relatives, forest elephants, remain poorly studied. Few decades ago, in the unique ‘natural enclosure experiment’ on Mount Cameroon, West/Central Africa, the rainforests were divided by lava flows which are not crossed by the local population of forest elephants. We assessed communities of trees, butterflies and two ecological guilds of moths in disturbed and undisturbed forests split by the longest lava flow at upland and montane elevations. Altogether, we surveyed 32 forest plots resulting in records of 2,025 trees of 97 species, and 7,853 butterflies and moths of 437 species. The disturbed forests differed in reduced tree density, height, and high canopy cover, and in increased DBH. Forest elephants also decreased tree species richness and altered their composition, probably by selective browsing and foraging. The elephant disturbance also increased species richness of butterflies and had various effects on species richness and composition of all studied insect groups. These changes were most probably caused by disturbance-driven alterations of (micro)habitats and species composition of trees. Moreover, the abandonment of forests by elephants led to local declines of range-restricted butterflies. Therefore, the current appalling decline of forest elephant populations across the Afrotropics most probably causes important changes in rainforest biodiversity and should be reflected by regional conservation authorities.

## 1. Introduction

Natural disturbances are key drivers of biodiversity in many terrestrial ecosystems (Grime, 1973; Connell, 1978), including tropical rainforests despite their traditional view as highly stable ecosystems (Chazdon, 2003; Burslem and Whitmore, 2006). Natural disturbances such as tree falls, fires, landslides, and insect herbivores outbreaks, generally open rainforest canopy, followed by temporarily changes microclimate and availability of plant resources (e.g., light, water, and soil nutrients) (Schnitzer *et al.*, 1991). The consequent changes in plant communities cause cascade effects on higher trophic levels (herbivores, predators, parasites), expanding the effects of disturbances on the entire ecosystem. Such increase of heterogeneity of habitats and species communities substantially contribute to maintaining the overall biodiversity of tropical forest ecosystems (Huston, 1979; Turner, 2010).

Megaherbivores, i.e. ≥1,000 kg herbivorous mammals, used to be among the main causes of such disturbances, before their abundances and diversity seriously dropped in all continents except Africa (Dirzo *et al.*, 2014; Galetti *et al.*, 2018). Among all megaherbivores, savanna elephants are best known to alter their habitats (e.g. Dirzo *et al.*, 2014; Guldemond *et al.*, 2017). Besides their important roles of seed dispersers or nutrient cyclers (Dirzo *et al.*, 2014), they directly impact savanna ecosystems through disturbing vegetation, especially by increasing tree mortality by browsing, trampling, and debarking (Guldemond *et al.*, 2017). Such habitat alterations substantially affect diversity of many organism groups (McCleery *et al.*, 2018), including insects. Savanna elephants were shown to positively influence diversity of grasshoppers (Samways and Kreuzinger, 2001) and dragonflies (Samways and Grant, 2008), whilst to have ambiguous effect on diversity of particular butterfly families (Bonnington *et al.*, 2008; Wilkerson *et al.*, 2013). Contrarily, too intensive disturbances caused by savanna elephants impact biodiversity negatively (O’Connor *et al.*, 2007; O’Connor and Page, 2014; Samways and Grant, 2008), similarly to other disturbance types.

Surprisingly, effects of forest elephants on biodiversity of Afrotropical rainforests remains strongly understudied (Guldemond *et al.*, 2017; Poulsen *et al.*, 2018). Although smaller (up to 5 tons, in comparison to 7 tons of savanna elephants), forest elephants are expected to affect their habitats by similar mechanisms as their savanna relatives, as recently reviewed by Poulsen *et al.* (2018). They were shown to impact rainforest tree density and diversity in both negative and positive ways (Campos-Arceiz and Blake, 2011; Hawthorne and Parren, 2000; Poulsen *et al.*, 2018). Besides local opening of forest canopy, they inhibit forest regeneration and maintain small-scaled canopy gaps (Omeja *et al.*, 2014, Terborgh *et al.*, 2016). However, the consequent cascade effects on rainforest biodiversity have not been studied yet, although rainforest organisms respond to other disturbances (Nyafwono, *et al.*, 2014; Alroy, 2017), and effects of elephant disturbances can be expected as well. Such research seems to be urgent especially because of the current steep decline of forest elephants across the Afrotropics (>60% decrease of abundance between 2002 and 2012; Maisels *et al.*, 2013). It has already resulted in local extinctions of forest elephants in numerous areas, including protected ones (Maisels *et al.*, 2013). In such situation, local policy makers and conservationists should be aware of potential changes in plant and animal communities to initiate more effective conservation planning.

In this study, we bring the first direct comparison of insect and tree communities in Afrotropical rainforests with and without forest elephants. Mount Cameroon provides an ideal opportunity for such study by offering a unique ‘natural enclosure experiment’. Rainforests on its southern slope were split by a continuous lava flow after eruptions in 1982 (from ca 1,400 m asl. up to ca 2,600 m asl., i.e. above the natural timberline) and 1999 (from the seashore up to ca 1,550 m asl.) (MINFOF, 2014; Fig. 1). Despite the slow natural succession on this lava flow, local forest elephants do not cross this barrier and stay on its western side close to three crater lakes, the only water sources during the dry seasons (MINFOF, 2014). Such unusual conditions represent a long-term enclosure experiment under natural conditions, performed on a much larger scale than any possible artificial enclosure study. In the disturbed and undisturbed sites, we sampled data on forest structure and communities of trees, butterflies, and two ecological groups of moths. We hypothesized that forest elephants changed the forest structure by opening its canopy, with the consequent changes in composition of all studied groups’ communities. We expected lower diversity of trees by the direct damage by elephants, and higher diversity of insects caused by the higher habitat heterogeneity. On the other hand, the ambiguous effect can be also hypothesized, as moths are known to be more closely dependent on diversity of trees (Beck *et al.*, 2002; Delabye *et al.*, in review), whilst butterflies rather benefit from canopy opening (Nyafwono *et al.*, 2015; Delabye *et al.*, in review). Finally, we focus on species’ distribution ranges in both types of forests, with no *a priori* hypothesis on the direction of the changes.

**Figure 1.**
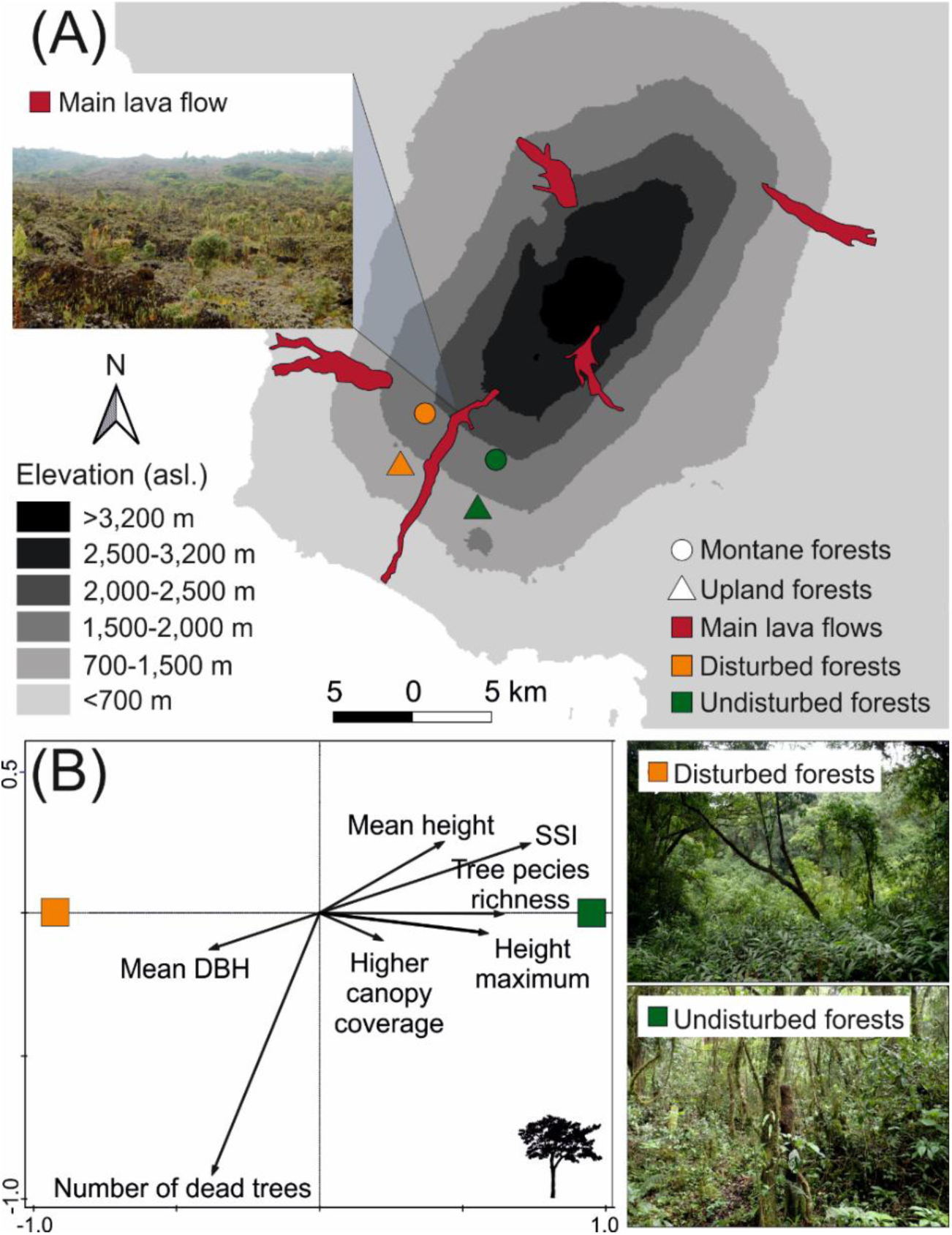
(A) Map of Mount Cameroon with the main lava flows and sampled forests. The pictures of disturbed and undisturbed forests were taken at the studied montane sites. (B) Redundancy analysis diagram visualizing effect of disturbance by elephants on forest structure.

## 2. Material and methods

### 2.1 Study area

Mount Cameroon (South-Western Province, Cameroon) is the highest mountain in West/Central Africa. This active volcano rises from the Gulf of Guinea seashore up to 4,095 m asl. Its southwestern slope represents the only complete altitudinal gradient from lowland up to the timberline (∼2,200 m asl.) of primary forests in the Afrotropics. Belonging to the biodiversity hotspot, Mount Cameroon harbor numerous endemics (e.g., Cable and Cheek, 1998; Ustjuzhanin *et al.*, 2018, 2020). With >12,000 mm of yearly precipitation, foothills of Mount Cameroon belong among the globally wettest places (Maicher *et al.*, 2020). Most of the rain falls during the wet season (June–September) with monthly precipitation >2,000 mm, whilst the dry season (late December–February) lacks any strong rains (Maicher *et al.*, 2020). Since 2009, most of its rainforests have become protected by the Mount Cameroon National Park.

Volcanism is the strongest natural disturbance on Mount Cameroon with frequent eruptions every ten to thirty years. Remarkably, on the studied southwestern slope, two eruptions in 1982 and 1999 created a continuous strip of bare lava rocks (hereinafter referred as ‘lava flow’) interrupting the rainforests on the southwestern slope from above the timberline down to the seashore (Fig. 1A).

A small population of forest elephants (*Loxodonta cyclotis*) strongly affect forests above ca. 800 m asl. on the southwestern slope (Cable and Cheek, 1998). It is highly isolated from nearest populations of the Korup National Park and the Banyang-Mbo Wildlife Sanctuary, as well as from much larger metapopulations in the Congo Basin (Blanc, 2008). It has been estimated to ∼130 individuals with a patchy local distribution (MINFOF, 2014). On the southwestern slope, they concentrate around three crater lakes representing the only available water sources during the high dry season (Ministry of Forestry and Wildlife of Cameroon, 2014). They rarely (if ever) cross the bare lava flows, representing natural obstacles dividing forests of the southwestern slope to two blocks with different dynamics. As a result, forests on the western side of the longest lava flow have an open structure, with numerous extensive clearings and pastures, whereas eastern forests are characteristic by undisturbed dense canopy (Fig. 1). Hereafter, we refer the forests west and east from the lava flow as *disturbed* and *undisturbed*, respectively. Effects of forest elephant disturbances on communities of trees and insects were investigated at four localities, two in an upland forest (1,100 m asl.), and two in a montane forest (1,850 m asl.).

### 2.3 Tree diversity and forest structure

At each of four sampling sites, eight circular plots (20 m radius, ∼150 m from each other) were established. All plots were established in high canopy forests (although sparse in the undisturbed sites), any larger clearings were avoided. In the disturbed forest sites, eight plots were selected to follow a linear transect among 16 plots previously used for a study of elevational diversity patterns (Hořák *et al.*, 2019; Maicher *et al.*, 2020), without looking at their diversity data. In the undisturbed forest sites, plots were established specifically for this study.

To assess the tree diversity in both elephant disturbed and undisturbed forest plots, all living and dead trees with diameter at breast height (DBH, 1.3 m) ≥10 cm were identified to (morpho)species (see Hořák *et al*. 2019 for details). To study impact of elephant disturbances on forest structure, each plot was also characterized by twelve descriptors. Besides *tree species richness, living* and *dead trees* with DBH ≥10 cm were counted. Consequently, DBH and basal area of each tree were measured and averaged per plot (*mean DBH* and *mean basal area*). Height of each tree was estimated and averaged per plot (*mean height*), together with the tallest tree height (*maximum height*) per plot. From these measurements, two additional indices were computed for each tree: stem slenderness index (SSI) was calculated as a ratio between tree height and DBH, and tree volume was estimated from the tree height and basal area (Poorter *et al.*, 2003). Both measurements were then averaged per plot (*mean SSI* and *mean tree volume*). Finally, following Grote (2003), proxies of shrub, lower canopy, and higher canopy coverages per plot were estimated by summing the DBH of three tree height categories: 0-8 m (shrubs), 8-16 m (lower canopy), >16 m (higher canopy).

### 2.4 Insect sampling

Butterflies and moths (Lepidoptera) were selected as the focal insect groups because they belong into one of the species richest insect orders, with relatively well-known ecology and resolved taxonomy, and with relatively well-standardized quantitative sampling methods. They also substantially differ in the usage of habitats, and together can be considered as useful biodiversity indicators. Within each sampling plot, fruit-feeding lepidopterans were sampled by five bait traps (four in understory, one in canopy) baited by fermented bananas (see Maicher *et al.*, 2020 for details). All fruit-feeding butterflies and moths (hereinafter referred as *butterflies* and *fruit-feeding moths*) were removed daily from the traps for ten consecutive days and identified to (morpho)species.

Additionally, moths were attracted by light at three ‘mothing plots’ per sampling site, established out of the sampling plots described above. These plots were selected to characterize the local heterogeneity of forest habitats and separated by a few hundred meters from each other. Moths were attracted by light (see Maicher *et al.*, 2020 for details) during six complete nights per elevation (i.e., two nights per plot). Six target moth groups (Lymantriinae, Notodontidae, Lasiocampidae, Sphingidae, Saturniidae, and Eupterotidae; hereafter referred as *light-attracted moths*) were collected manually and identified into (morpho)species. The three lepidopteran datasets (butterflies, and fruit-feeding and light-attracted moths) were extracted from Maicher *et al.* (2020) for the disturbed forest plots, whilst the described sampling was performed in the undisturbed forest plots specifically for this study. Voucher specimens are deposited in the Nature Education Centre, Jagellonian University, Kraków, Poland.

To partially cover the seasonality (Maicher *et al.*, 2018), the insect sampling was repeated during transition from wet to dry season (November/December), and transition from dry to wet season (April/May) in all disturbed and undisturbed forest plots.

### 2.5 Diversity analyses

To check sampling completeness of all focal groups, the sampling coverages were computed to evaluate our data quality using the *iNEXT* package (Hsieh *et al.*, 2019) in R 3.5.1 (R Core Team, 2018). For all focal groups in all seasons and at all elevations, the sampling coverages were always ≥0.84 (mostly even ≥0.90), indicating a sufficient coverage of the sampled communities (Table S1). Therefore, observed species richness was used in all analyses (Beck & Schwanghart, 2010).

Effects of *disturbance* on species richness were analyzed separately for each focal group by Generalized Estimated Equations (GEE) using the *geepack* package (Højsgaard *et al.*, 2006). For trees, species richness from individual plots were used as a ‘sample’ with an independent covariance structure, with *disturbance, elevation*, and their interaction treated as explanatory variables. For lepidopterans, because of the temporal pseudo-replicative sampling design, species richness from a sampling day (butterflies and fruit-feeding moths) or night (light-attracted moths) at individual plot was used as a ‘sample’ with the first-order autoregressive relationship *AR(1)* covariance structure (i.e. repeated measurements design). *Disturbance, season, elevation, disturbance*season*, and *disturbance*elevation* were treated as explanatory variables. All models were conducted with Poisson distribution and log-link function. Pairwise post-hoc comparisons of the estimated marginal means were compared by Wald χ^2^ tests.

Differences in composition of communities between the disturbed and undisturbed forests were analyzed by multivariate ordination methods (Šmilauer & Lepš, 2014), separately for each focal group. Firstly, the main patterns in species composition of individual plots were visualized by Non-Metric Multidimensional Scaling (NMDS) in Primer-E v6 (Clarke & Gorley, 2006). NMDSs were generated using Bray-Curtis similarity, computed from square-root transformed species abundances per plot. Subsequently, influence of *disturbance* on community composition of each focal group was tested by constrained partial Canonical Correspondence Analyses (CCA) with log-transformed species’ abundances as response variables and *elevation* as covariate (Šmilauer & Lepš, 2014). Significance of all partial CCAs were tested by Monte Carlo permutation tests with 9,999 permutations.

Finally, differences in the forest structure descriptors between the disturbed and undisturbed forests were analyzed by partial Redundancy Analysis (RDA). Prior to the analysis, preliminary checking of the multicollinearity table among the structure descriptors was investigated. Only forest structure descriptors with pairwise collinearity <0.80, i.e. tree species richness, number of dead trees, mean DBH, mean height, maximum height, mean SSI, and higher canopy coverage, were included in these analyses. Their log-transformed values were used as response variables (Šmilauer & Lepš, 2014). RDA was then run with *disturbance* as explanatory variable and *elevation* as covariate, and tested by Monte Carlo permutation test (9,999 permutations). All CCAs and RDA were performed in Canoco 5 (ter Braak & Šmilauer, 2012).

### 2.6 Species distribution range

To analyze if the elephant disturbance supports rather range-restricted species or widely distributed generalists, we used numbers of Afrotropical countries with known records of each tree and lepidopteran species as a proxy for their distribution range; we are not aware of any more precise existing dataset covering all studied groups for the generally understudied Afrotropics. Because of the limited knowledge on Afrotropical Lepidoptera, we ranked only butterflies and light-attracted Sphingidae and Saturniidae moths (the latter two analyzed together and referred as *light-attracted moths*). This distribution data were excerpted from the RAINBIO database for trees (Dauby *et al.*, 2016), Williams (2018) for butterflies, and Afromoths.net for moths (De Prins & De Prins, 2018); all considered as the most comprehensive databases. Non-native tree species and all morphospecies were excluded from these analyses. In total, 73 species of trees, 71 butterflies, and 21 moths were included in the distribution range analyses.

To consider the relative abundances of individual species in the communities, the distribution range of each species was multiplied by the number of collected individuals per sample and their sums were divided by the total number of individuals recorded at each sample. These *mean distribution ranges* per sample were then compared between disturbed and undisturbed forest sites by GEE analyses (with normal distribution; independent covariance structure) following the same model design as for the above-described comparisons of species richness.

## 3. Results

In total, 2,025 trees were identified to 97 species and 7,853 butterflies and moths were identified to 437 species in all sampled forest plots (Table S1).

### 3.1 Elephant disturbances and structure of rainforests

The partial-RDA ordination analysis showed significant differences in the forest structure descriptors between the disturbed and undisturbed forests (Fig. 1B). In total, the two main ordination axes explained 18.5% of the adjusted variation (all axes eigenvalues: 0.83; Pseudo-F = 7.8; p = 0.002). In the disturbed plots, *tree species richness, mean SSI, mean height, maximum height*, and *higher canopy coverage* were lower. In contrast, *mean DBH* was larger in the disturbed forests (Fig. 1B).

### 3.2. Elephant disturbances and tree diversity

Elephant disturbances affected tree species richness per sampled elevation, as well as per sampled plot. In both upland and montane forests, total tree species richness of the disturbed sites was nearly half in comparison to the undisturbed sites (Fig. 2A; Table S1). Tree species richness per plot was significantly affected by disturbance (higher at undisturbed forest plots) and elevation (higher at the upland forests) (Fig. 2B; Table 1A).

**Table 1.**
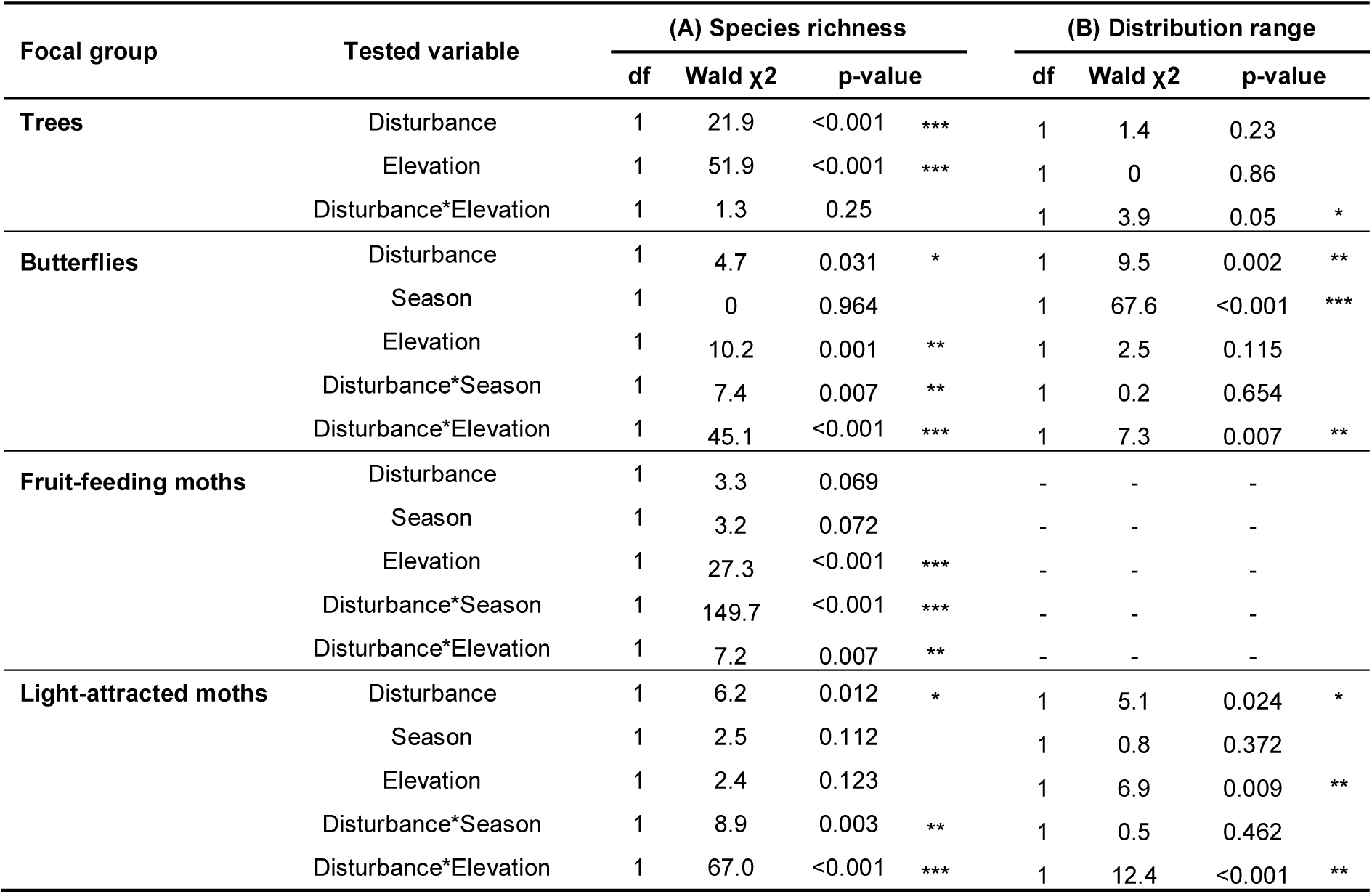
Results of the GEE models comparing (A) tree and insect species richness per plot in forests disturbed and undisturbed by elephants (with included effects of elevation, season, and their interactions into the models), and (B) mean distribution range of trees and insects per plot (*p <0.05; **p <0.01; ***p <0.001). See methods for the model details.

**Figure 2.**
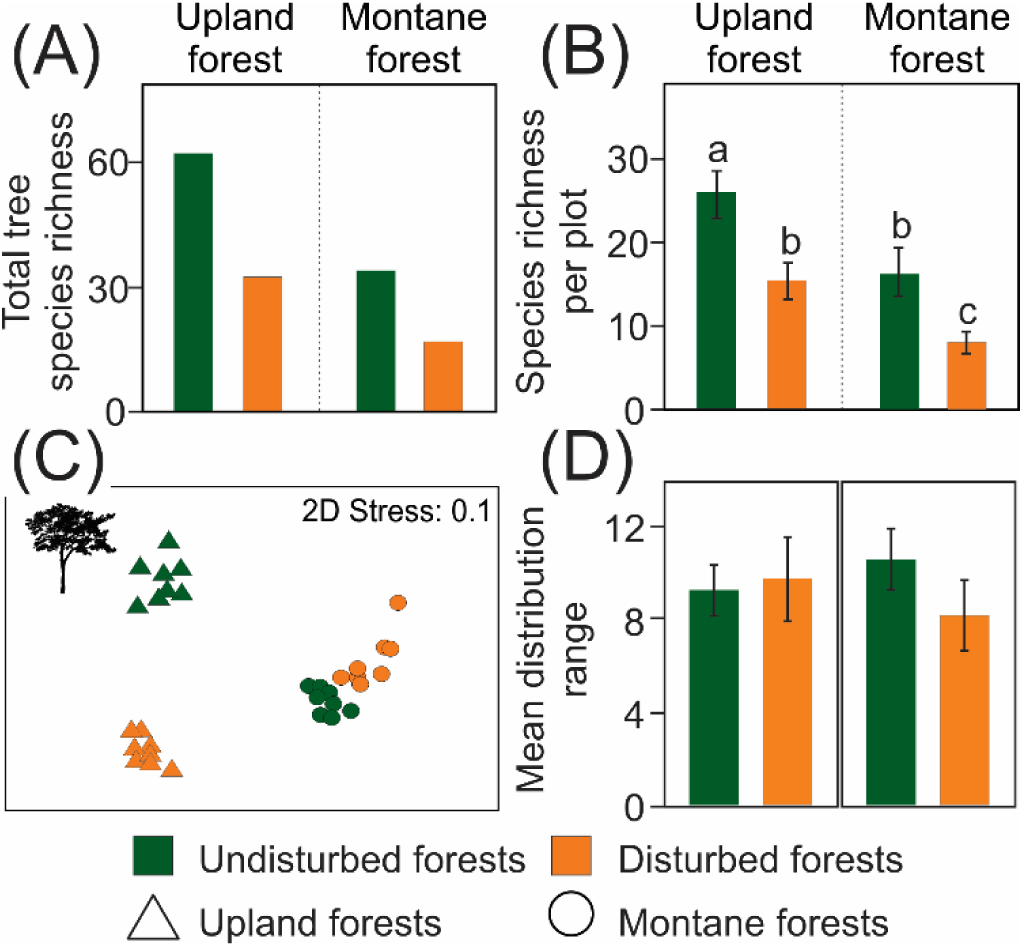
Differences in tree species richness, community composition, and mean distribution range between forests disturbed and undisturbed by elephants. Tree species richness per (A) forest site, and (B) per sampling plot estimated by GEE (estimated means with 95% unconditional confidence intervals). The letters visualize results of the post-hoc pairwise comparisons. (C) NMDS diagrams of the tree community compositions at the sampled forest plots. (D) Mean distribution range of trees per sampling plot estimated by GEE (estimated means with 95% unconditional confidence intervals).

Tree communities significantly differed in composition between the forests disturbed and undisturbed by elephants according to the partial-CCA (all-axes eigenvalues: 4.55; Pseudo-F = 3.8; p < 0.001). The first NMDS axis reflected elevation, whilst the tree communities of the disturbed and undisturbed forests were relatively well-separated along the second axis (Fig. 2C). The ordination diagram also showed relatively higher dissimilarities of tree communities between the disturbed and undisturbed plots at the upland than at the montane forests (Fig. 2C).

### 3.3. Elephant disturbances and insect diversity

The responses of individual insect groups’ total species richness per sampling site to elephant disturbances were rather inconsistent among the studied elevations and seasons. Butterflies and fruit-feeding moths showed lower total species richness in the disturbed forests at both elevations during the transition from wet to dry seasons, which became higher or comparable to the undisturbed forests during the transition from dry to wet seasons (Fig. 3a,b; Table S1). Light-attracted moths were species-richer in the disturbed upland forest than in the undisturbed upland forest during both sampled seasons but species-poorer in the montane forest during both sampled seasons (Fig. 3c; Table S1).

**Figure 3.**
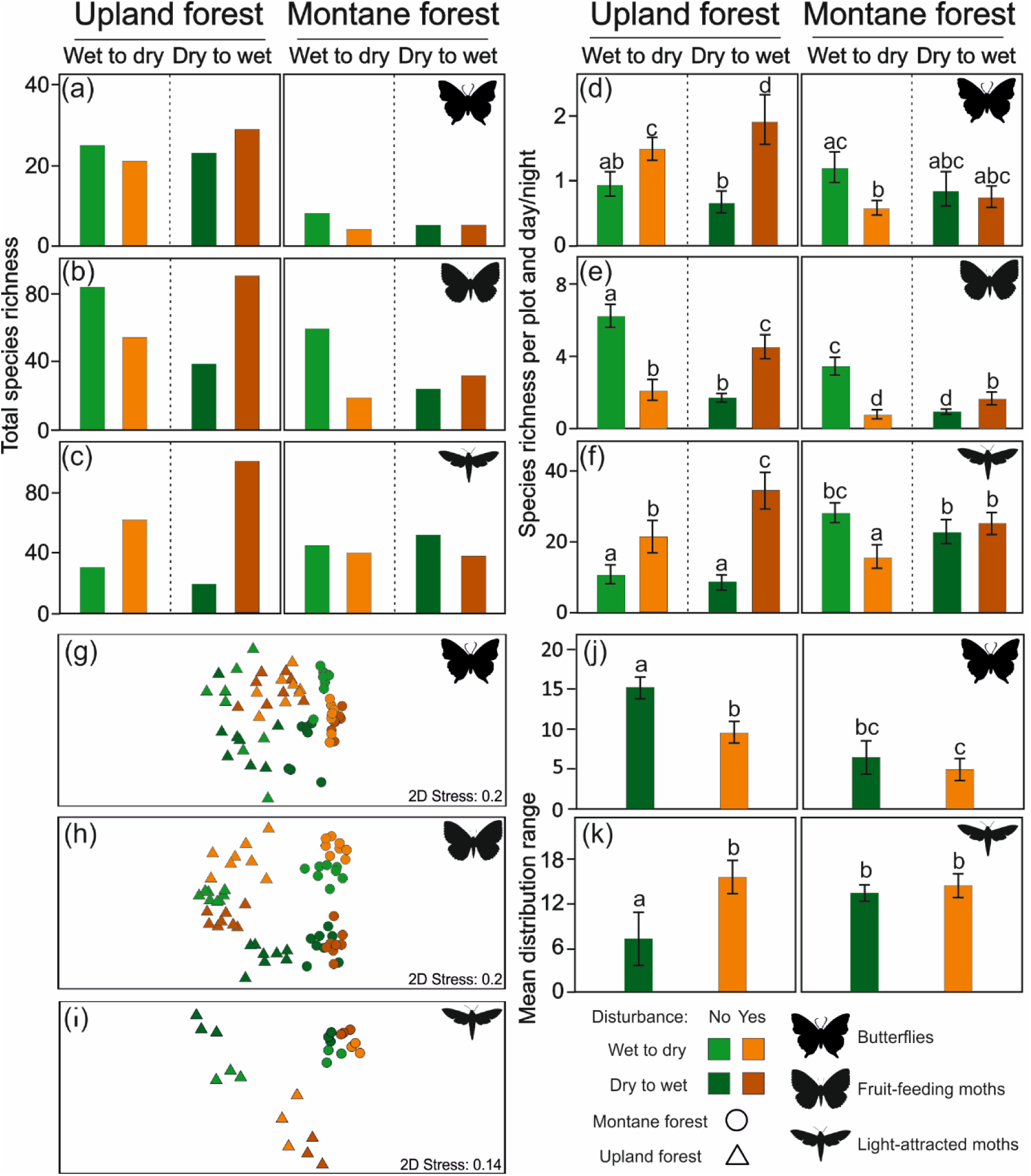
Species richness of insects per sampling site and season (a-c), and sampling plots and day or night (d-f) as estimated by GEEs (estimated means with 95% unconditional confidence intervals are visualized). (g-i) NMDS diagrams of insect community compositions at the sampled forest plots. (j, k) Mean distribution range of insects estimated by GEEs. Letters visualize results of the post-hoc pairwise comparisons.

The effects of elephant disturbances on insect species richness per plot also differed among the studied insect groups. The interactions disturbance*season and disturbance*elevation were significant for all insect groups (Table 1), indicating complex effects of elephant disturbances on insect species richness. GEEs showed a significant positive effect of elephant disturbances on species richness of butterflies and light-attracted moths (Fig. 3d,f; Table 1). No significant effect of elephant disturbances was detected for fruit-feeding moths (Table 1). Both butterflies and fruit-feeding moths were significantly species richer at the lower altitudes, whilst no significant effect of elevation on light-attracted moths was revealed (Fig. 3d-f; Table 1). Insignificant effects of season were shown for all studied insect groups (Table 1). For butterflies and light-attracted moths, the pairwise post-hoc comparisons of disturbed and undisturbed forests showed that species richness was significantly higher in the disturbed upland forests for both groups, and significantly lower or not significantly different (depending on the sampled season) in the montane forests (Fig. 3d,f). In contrast, fruit-feeding moth species richness was significantly lower in the disturbed forests at both elevations during the transition from wet to dry season, but significantly richer during the transition from dry to wet season (Fig. 3e).

Elephant disturbances significantly affected species composition of all focal insect groups in partial CCAs (butterflies: all-axes eigenvalue: 2.75; Pseudo-F: 4.6; p-value: <0.001; fruit-feeding moths: all-axes eigenvalue: 5.27; Pseudo-F: 3.2; p-value: <0.001; light-attracted moths: all-axes eigenvalue: 2.96; Pseudo-F: 4.5; p-value: <0.001). For butterflies and fruit-feeding moths, the first NMDS axes can be related to elevation, in contrast to light-attracted moths where elevation can be related to the second NMDS axis (Fig. 3g-i). All groups were well-clustered according to the disturbance type at both elevations. The effect of disturbance was interacting with season and elevation for all groups (Fig. 3g-i). Among all insect groups, light-attracted moths species composition responded to elephant disturbances very similarly to trees, with well-separated upland disturbed and undisturbed forest types and comparatively less heterogenous montane forest samples (Fig. 2C; Fig. 3i).

### 3.4 Elephant disturbances and species’ distribution range

Elephant disturbances and elevation showed marginally significant effects of their interaction on distribution range of tree species, although no significant separate effect was detected for them (Table 1B). In the undisturbed forests, the mean tree species’ distribution range was positively associated with increasing elevation, while negatively associated with increasing elevation in the disturbed forests. However, the pairwise post-hoc comparisons were insignificant (Fig. 2D).

Patterns of distribution range differed between the two analyzed insect groups. Butterfly species’ distribution range was significantly lower at high elevation and in the disturbed forests (Fig. 3j). Similarly, moths’ mean distribution range was significantly affected by elephant disturbances and seasons (Table 1B). Nevertheless, pairwise post-hoc comparisons showed that light-attracted moths in the undisturbed upland forest had a significantly lower distribution range than in all other studied forests, which did not significantly differ from each other (Fig. 3k).

## 4. Discussion

Our study has shown a strong effect of forest elephants on rainforest biodiversity. Concordant to our first hypothesis, their long-term absence at the studied forests changed the forest structure. It has led to an increase of forest height, closure of its canopy, and dominance of smaller over large trees. This observed shift in rainforest structure can be interpreted by a combination of direct and indirect effects driven by forest elephants. Because of their high appetite and large body size, forest elephants surely eliminate some trees (Terborgh *et al.*, 2016). They directly consume high amount of tree biomass, as well as their fruits and seeds (Blake, 2002). When struggling through forest, elephants break stems and sometimes even uproot trees, while their repeated trampling denude the forest floor and destroy fallen seeds and saplings (Terborgh *et al.*, 2016). Moreover, the direct damages are likely to increase tree susceptibility to pathogens. Although the number of dead trees seemed to poorly characterize the disturbed forests, the higher tree density in the undisturbed plots supports this hypothesis. Thus, the presence of a few large trees in the plots disturbed by forest elephants can be explained by only a small portions of trees escaping the browsing pressure (Terborgh *et al.*, 2016).

Together with altering the rainforest structure, forest elephants decreased tree species richness and change tree community composition, confirming our second hypothesis. Although forest elephants are generalized herbivores (Blake, 2002), they prefer particular species of trees and other plants (Blake, 2002). Thereby, their selective browsing of palatable species affects tree mortality and recruitment, which can explain the observed differences in tree communities between the disturbed and undisturbed forests. Finally, similarly as in savanna, we can reasonably expect different resistance of tree species to repeated disturbances by forest elephants, or differences in their ability to recover from damages (Owen-Smith *et al.*, 2019). Unfortunately, the knowledge of African forest elephants’ browsing preferences and/or Afrotropical trees’ resistance to disturbances are not enough to decide which effect prevails in the alterations of rainforest structure by elephants.

The presence of forest elephants impacted all studied herbivorous insect communities as well, although differently for particular insect groups. These can be related to the changes in composition of tree communities and in habitat structure in the disturbed forests. The upland rainforests disturbed by elephants harbored more species of butterflies and light-attracted moths. However, all other effects of disturbances differed according the studied elevation and season, as well as among the insect groups. Tropical butterflies rely on forests gaps and solar radiation for their thermoregulation (Clench, 1966) and oviposition on larval food-plants (mostly herbs; Hill *et al.*, 2001), therefore their diversity decrease after the upland forest elephants enclosure cannot be surprising. By opening of rainforest canopy, forest elephants seem to support quantity and heterogeneity of resources available for butterflies (Delabye *et al.*, in review). However, such hypothesis can hardly explain the detected decrease of light-attracted (night flying) moth diversity in the undisturbed upland rainforests. In fact, diversity of moths has been repeatedly shown to increase with diversity of trees, as the most common food plants for their caterpillars (Janzen, 1988; Tews *et al.*, 2004). Therefore, the opposite effect of disturbance by forest elephants can be expected. Unfortunately, we do not have any other explanation of the positive effect of forest disturbances in the sampled upland forests. Contrastingly, fruit-feeding moths are relatively independent to forest structure (Delabye *et al.*, unpublished). They can follow the spatiotemporal changes of ripe fruits (adult food) or young sprouts (larval food) more tightly than fruit-feeding butterflies, which could partly explain their seasonally inconsistent reaction to the elephant disturbances. Unfortunately, no data to confirm or reject such hypothesis exist.

In the montane forests, we have not found any consistent changes of the insects’ diversity, as it strongly varied with season and studied insect group. Moreover, the communities of all insect groups were highly homogeneous in both forest types in this high elevation. The montane forests on Mount Cameroon are already relatively open and with limited tree diversity (Hořák *et al.*, 2019) that additional disturbances by elephants could hardly increase habitat heterogeneity even for butterflies. Moreover, some tree dominants in the montane forests, such as *Schefflera abyssinica* and *S. mannii*, are (semi)deciduous during the dry season which generally open the higher canopy even in the undisturbed forests. Simultaneously, these dominants get typically recruited as epiphytes, later strangling their hosts (Abiyu *et al.*, 2013). Therefore, they may more efficiently escape from any elephant effects. We hypothesize that these effects together result in more similarity between the disturbed and undisturbed forests at higher elevations. Last but not least, we have recently revealed a strong seasonal shift in elevational ranges of both butterflies and moths (Maicher *et al*., 2020); the seasonal discrepancies in the effect of disturbance could be related to it. Unfortunately, we do not have any detailed data on this phenomenon from the undisturbed forest plots.

Recently, Poulsen *et al.* (2018) discussed the fate of Afrotropical rainforests of a future world without forest elephants. The authors hypothesized that their loss would increase understory stem density and change tree species composition. We concur with Poulsen’s hypotheses from our data study. Moreover, we have shown that the change of forest structure and composition can have strong cascading effects on other trophic levels, at least in the upland rainforests. Hawthorne and Parren (2000) demonstrated that the disappearance of forest elephants from several Ghanaian forests did not have any remarkable effect on plant populations at the country level. However, our study has shown that the local consequences of forest elephants’ disappearance can be highly significant for trees, as well as for higher trophic levels. Although more comparative studies are required, forest elephant extinction would accelerate the vegetation succession, enclose the rainforest canopy, and generally impoverish the habitat heterogeneity in Afrotropical rainforests. These would be unavoidably followed by changes in rainforest communities and by declines of range-restricted species that profit from disturbances, as we have shown for some of the herbivorous insects in the upland rainforests.

In conclusion, our study showed that African forest elephants contribute for maintaining the rainforest heterogeneity and tree diversity. The elephant-related habitat heterogeneity increased the heterogeneity of available niches and sustain diverse communities of Afrotropical insects. Despite the lack of any data, we can even speculate on the consequences on biodiversity at other trophic levels. Therefore, we have confirmed the African forest elephant as a key-stone species in the Afrotropical rainforest ecosystems. The maintenance of forest elephant populations in Afrotropical rainforests appears to be necessary to prevent biodiversity declines. Unfortunately, the decline of forest elephant populations in West and Central African rainforests is alarming, and most probably would be followed by other species extinctions. It is even highly probable that such processes are already ongoing, although unrecorded in one of the least studied biogeographic areas in the world. Therefore, we urge for more efficient conservation of the remaining populations of forest elephants. Their effects on the entire rainforest ecosystems must be recognized and incorporated into the management plans of Afrotropical protected areas.

## Acknowledgments

We are grateful to Francis E. Luma, Nestor T. Fominka, Jacques E. Chi, Congo S. Kulu, and other field assistants for their help in the field; Štěpán Janeček, Szabolcs Sáfián, Jan E.J. Mertens, Jennifer T. Kimbeng, and Pavel Potocký for help with Lepidoptera sampling at the elephant-disturbed plots; Karolina Sroka, Ewelina Sroka, and Jadwiga Lorenc-Brudecka for Lepidoptera setting; Elias Ndive for tree identification; Yannick Klomberg for reviewing distribution of trees; Axel Hausmann for access to the Bavarian State Collection of Zoology; and the Mount Cameroon National Park staff for their support. This study was performed under authorizations of the Cameroonian Ministries for Forestry and Wildlife, and for Scientific Research and Innovation. Our project was funded by Czech Science Foundation (16-11164Y, 17-19376S), University of South Bohemia (GAJU 030/2016/P, 038/2019/P), Charles University (PRIMUS/17/SCI/8, UNCE204069), and Czech Academy of Sciences (RVO 67985939).

## Data availability

Data available via the Zenodo repository (*doi will be provided after acceptance*).

