## Supplementary material for "Does rainforest biodiversity stand on the shoulders of giants? Effect of disturbances by forest elephants on trees and insects on Mount Cameroon": Table S1

**APPENDIX S1.** Summary of abundance and diversity of trees and insects in forests disturbed and undisturbed by elephants on Mount Cameroon.

**Table S1.** Abundance and diversity of trees and insects in different seasons in forests disturbed and undisturbed by elephants.

|  |  | **Undisturbed forests** | | | | |  | **Disturbed forest** | | | | |
| --- | --- | --- | --- | --- | --- | --- | --- | --- | --- | --- | --- | --- |
|  |  | **1,100 m a.s.l.** | |  | **1,850 m a.s.l.** | |  | **1,100 m a.s.l.** | |  | **1,850 m a.s.l.** | |
|  |  | **Wet to dry** | **Dry to wet** |  | **Wet to dry** | **Dry to wet** |  | **Wet to dry** | **Dry to wet** |  | **Wet to dry** | **Dry to wet** |
| **Trees** | **Abundance** | 802 | |  | 438 | |  | 511 | |  | 274 | |
|  | **Species richness** | 62 | |  | 32 | |  | 32 | |  | 16 | |
|  | **Sampling coverage** | 0.98 | |  | 0.99 | |  | 0.99 | |  | 0.99 | |
| **Butterflies** | **Abundance** | 74 | 67 |  | 355 | 95 |  | 255 | 193 |  | 68 | 119 |
|  | **Species richness** | 25 | 23 |  | 8 | 5 |  | 21 | 29 |  | 4 | 5 |
|  | **Sampling coverage** | 0.88 | 0.85 |  | 0.99 | 0.99 |  | 0.97 | 0.95 |  | 1.00 | 0.99 |
| **Fruit-feeding moths** | **Abundance** | 1,806 | 184 |  | 458 | 93 |  | 192 | 499 |  | 101 | 144 |
|  | **Species richness** | 85 | 39 |  | 60 | 24 |  | 55 | 92 |  | 19 | 32 |
|  | **Sampling coverage** | 0.98 | 0.91 |  | 0.94 | 0.85 |  | 0.86 | 0.92 |  | 0.94 | 0.85 |
| **Light-attracted moths** | **Abundance** | 326 | 61 |  | 633 | 473 |  | 208 | 469 |  | 383 | 597 |
|  | **Species richness** | 30 | 19 |  | 45 | 52 |  | 62 | 101 |  | 40 | 38 |
|  | **Sampling coverage** | 0.97 | 0.84 |  | 0.98 | 0.96 |  | 0.86 | 0.90 |  | 0.96 | 0.99 |
